# Comparing Alternative Computational Models of the Stroop Task Using Effective Connectivity Analysis of fMRI Data

**DOI:** 10.1101/647271

**Authors:** Micah Ketola, Linxing Preston Jiang, Andrea Stocco

## Abstract

Methodological advances have made it possible to generate fMRI predictions for cognitive architectures, such as ACT-R, thus expanding the range of model predictions and making it possible to distinguish between alternative models that produce otherwise identical behavioral patterns. However, for tasks associated with relatively brief response times, fMRI predictions are often not sufficient to compare alternative models. In this paper, we outline a method based on *effective connectivity*, which significantly augments the amount of information that can be extracted from fMRI data to distinguish between models. We show the application of this method in the case of two competing ACT-R models of the Stroop task. Although the models make, predictably, identical behavioral and BOLD time-course predictions, patterns of functional connectivity favor one model over the other. Finally, we show that the same data suggests directions in which both models should be revised.

## Introduction

One of the traditional problems in the field of cognitive modeling is deciding which of two alternative models provides best explains a phenomenon. Traditionally, the most common approach has been to compare models using null-hypothesis testing procedures. In essence, conditions are identified in which the two models make qualitatively different predictions, and the hypothesized pattern is tested using classical statistical testing techniques. While more sophisticated approaches have been proposed (Pitt, Kim, Navarro, & Myung, 2006), this approach remains the *de facto* standard of the field.

The search for conditions in which two models differ is sometimes strenuous, as the same external behavior can occasionally be obtained through different possible internal processes and model parameters. By shedding light on more direct correlates of cognitive processes, neuroimaging data provides a potential way to distinguish between otherwise behaviorally identical results (Sohn et al., 2004). For this reason, procedures have been devised to derive neuroimaging predictions from computational models, most commonly in the domain of fMRI (Anderson, Fincham, Qin, & Stocco, 2008; Borst, Nijboer, Taatgen, van Rijn, & Anderson, 2015).

While the use of fMRI has greatly expanded upon the possible predictions that can distinguish between the two models, a number of limitations still exist. A main limitation arises from poor temporal resolution of fMRI. The BOLD signal that is recorded in MRI scanners is extremely sluggish, and peaks approximately five seconds after an event. This poses a problem for resolving cognitive processes that occur quickly in time.

Other neuroimaging methods, such as EEG and MEG, offer much greater temporal resolution, but they trade off this advantage with much lower spatial resolution. Furthermore, the oscillatory nature of EEG and MEG signals further complicates the process of deriving predictions from models, as changes in raw signals can occur at different frequency bands (van Vugt, 2014).

Even if these technical issues could be solved, a deeper problem is that the most common methods devised to compare models against neuroimaging data focus on accounting for the common time course of brain activity and model computations. But models, by their very nature, usually make richer predictions about the internal dynamics that lead to either brain activity or behavioral responses. For example, models often make specific assumptions about the *directionality* of an effect, or about how different model components interact with each other. These predictions cannot be tested by simply correlating neuroimaging time series with the order of computations.

In this paper, we describe and demonstrate an alternative and novel method to test models using neuroimaging data. This method is based on patterns of *effective connectivity* between brain regions. “Effective connectivity” is an umbrella term to characterize the functional exchange of information between two brain regions, based on the analysis of their respective time series. Because effective connectivity provides measures of directional communication between two regions, it can be used to examine the internal dynamics of a computational model. Furthermore, because effective connectivity can be estimated from either fMRI or EEG data, it expands the dimensions across which models can be compared without requiring collecting additional data.

In the remainder of this paper, we will outline our method and apply it to a specific, and exquisitely cognitive case, namely, determining which of two prominent computational explanations for the Stroop interference best explains the data.

### ACT-R

Although our method could be applied to any computational model, for convenience, it will be demonstrated with two models developed in the Adaptive Control of Thought–Rational (ACT-R) cognitive architecture (Anderson et al., 2004). This choice was made for three reasons. First, ACT-R is the most successful and widespread architecture, having been used in hundreds of publications since its inception, and by far the most popular in the field of cognitive research (Kotseruba & Tsotsos, 2018); Thus, it provides an excellent domain in which to demonstrate the procedure. Second, ACT-R already provides well-tested mappings between architectural components and brain regions with established procedures to predict fMRI activity from model simulations. Therefore, these assumptions can be adopted without the need to provide additional justifications. Finally, the assumptions of ACT-R provide a reasonable mechanism to translate model activity into effective connectivity. As it will be shown, this is based on the functional requirements of the procedural module, which have been examined and discussed in the past.

ACT-R represents knowledge in two formats, *declarative* and *procedural*. Declarative knowledge is made of record-like structures, called *chunks*, which capture semantic memories, perceptual inputs, and motor commands. Procedural knowledge consists of production rules (or simply “productions”), state-action pairs that encode the specific policy to perform a task. In summary, chunks represent information, and productions act upon them.

Chunks are processed by functionally specialized modules. For instance, perceptual modules create new chunks to represent the contents of the outside world, and a memory module maintains chunks in long-term memory. Each module contains one or more buffers, limited-capacity stores that contain at most one chunk. Buffers are the only mechanisms through which chunks and productions interact: Chunks can be inspected, copied, and modified by productions when exposed into buffers.

As noted above, much work has been dedicated to map ACT-R modules to corresponding neural circuits. This work has yielded a number of reliable functional mappings, including the association between anterior cingulate cortex and the goal buffer in the goal module, between the lateral prefrontal cortex and the retrieval buffer of the long-term memory module, between posterior parietal cortex and the imaginal buffer of working memory, between the fusiform gyrus and the visual buffer in the visual module, and between the primary motor cortex and the manual buffer in the motor module (Fincham & Anderson, 2006; Sohn, Albert, Jung, Carter, & Anderson, 2007; Danker, Gunn, & Anderson, 2008; Anderson et al., 2004, 2008). These five modules will be the focus of this paper.

### Dynamic Causal Modeling

To estimate effective connectivity, we adopted a framework known as Dynamic Causal Modeling (DCM) (Friston, Harrison, & Penny, 2003). In essence, DCM is procedure to model the time-course of in brain activity in a set of brain regions through a dynamical system of other brain regions and event vectors. Specifically, the time course of activity of a region *i* is expressed as a bilinear state equation:

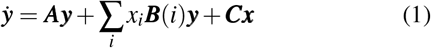

where ***y*** are the time series of neuronal activities and ***x*** are the time series of the events. ***A*** defines intrinsic connectivity between different regions (fixed connectivity), ***C*** defines effects by task inputs, and ***B*** defines the modular effects that task conditions have on the connectivity between regions (modulation of connectivity).

### ACT-R Predictions for Effective Connectivity

Because effective connectivity can be interpreted as directional effects between cortical regions, a direct link can be made between this measure and the nature of ACT-R computations. As discussed above, ACT-R works by firing one production at a time during its cognitive cycle; this production, in turn, changes the state of the system by modifying or copying information from one buffer to the other. For example, in what is perhaps the most common operation in ACT-R models, a production rule extracts values from the slots of chunks placed in either the imaginal or the visual buffer (to extract contextual task information) and places them in the retrieval buffer, so that they function as cues for retrieving relevant information from long-term memory. In fact, production rules are the only way information is exchanged between modules.

Given their role in coordinating module-to-module communication, we made the assumption that patterns of effective connectivity can be derived by the analysis of information transferred carried by out in the sequence of production rules firing.

On the surface, this idea runs against the established identification between production rules (and their associated procedural module) and the activity of the basal ganglia (Anderson et al., 2004; Anderson, 2007). The two interpretations, however, are not incompatible with each other. Anderson et al. (2008) had previously suggested that common functional connectivity patterns in the brain reflect the ubiquity of common operations that exchange information between different buffers; the example production given above is one of those put forward by the authors. It has also been noted before that the function of the basal ganglia is to direct inputs to cortical regions, a role that is both compatible with the procedural module and with the proposed interpretation of effective connectivity (Stocco, Lebiere, & Anderson, 2010). Finally, a recent study that combined ACT-R modeling and Transcranial Magnetic Stimulation (Rice & Stocco, 2019) has provided evidence that production rules do not only reflect the activity of the basal ganglia but also, more generally, the direct exchange of information between cortical regions. Thus, we believe that the hypothesis that production rules could be used to estimate effective connective is a plausible one.

In this study, the relationship between production rules and effective connectivity was operationalized in the following, simple algorithm. First, an *N* × *N* squared matrix ***E***, with *N* being the number of buffers examined, is generated and initialized to zeros. Then, the target model is run and its trace is segmented into epochs of interest (e.g., all the trials of the same conditions). The structure of each production rule firing within that epoch is then examined. For each variable in the production rule, the *source* buffer *S* at which the variable is introduced (or, technically, bound to a value) in the left-hand side and the *target* buffer *T* in which the bound value is placed are recorded. The value of the matrix cell ***E***_*S,T*_ is then incremented by one. If a variable appears in multiple source buffers *S*_1_, *S*_2_ … *S*_*N*_ or target buffers *T*_1_, *T*_2_ … *T*_*N*_, then all the cells ***E***_*i*∈*N,j*∈*N*_ are updated. When all the productions have been examined, ***E*** is taken to represent the predicted effective connectivity for that particular condition.

## An Application of the Method: ACT-R Models of the Stroop Task

This method was demonstrated using two competing models of the Stroop task. In the Stroop task, participants are shown a colored character string and asked to report the color of the character string. The character string can either be congruent with the color (“RED” printed in red), incongruent (“BLUE” printed in green), or neutral (“CHAIR” printed in blue). The typical finding is that reaction times in each condition are significantly different from one another, with congruent trials being the fastest, incongruent trials being the slowest, and neutral trials in between (Bugg, McDaniel, Scullin, & Braver, 2011). This difference in reaction times between trial types is referred to as Stroop interference.

The two models were adapted versions of two previously proposed models of the Stroop task, authored by Lovett (2005) and by Altmann and Davidson (2001), respectively. Since both models were published before ACT-R was modified to account for neuroimaging data, they had to be reimplemented in the most recent version of ACT-R (version 7.6). This processes also ensured that the two models interacted with the task using the same sensorimotor mechanisms, i.e. visual objects and responses were given in the same way. From now on, we will refer to these two models as the Altmann-like model and the Lovett-like model.

The re-implemented models maintained the underlying assumptions of their original versions. Specifically, the two models provide different explanations about the nature of Stroop interference. In the Altmann model (Figure 1A), Stroop interference is driven by interference at the lemma layer. When a word is processed, it has direct access to its’ lemma, or conceptual representation. Access to the lemma of a color is indirect, requiring an extra retrieval not seen with words. The model assumes that the word dimension of the Stroop stimulus is automatically processed first, therefore activating the lemma attached to the word dimension of the stimulus. As it tries to process the color dimension of the stimulus, the word-lemma is active and can either facilitate or inhibit retrieval of the correct color-lemma. In cases of facilitation, activation from the word-lemma spreads to the coinciding color-lemma, increasing the likelihood of correct retrieval on congruent trials. Oppositely, on incongruent trials, this activation spreads to the incorrect color-lemma, creating increased competition between color-lemmas and introducing ambiguity. For neutral trials, the word-lemma has no corresponding color-lemma, resulting in neither facilitation nor inhibition. The color-lemma is compared to visual cues and re-selected if inconsistent or otherwise used in further processing. A manual response is then retrieved using the color-lemma, and used to press a key on the keyboard.

**Figure 1:**
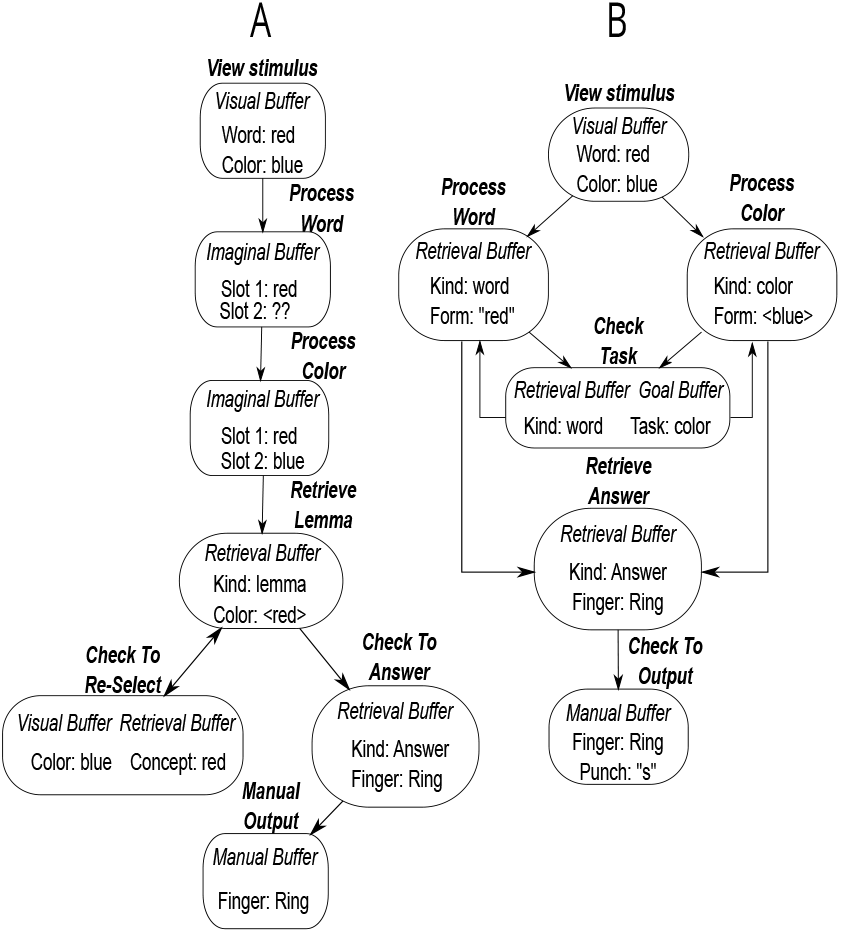
Flow-chart representation of the strategies used by the Altmann model (“A”, left) and the Lovett model (“B”, right) when processing Stroop trials.

In the Lovett model (Figure 1B), Stroop interference is driven by the competition between alternative *word-association chunks*, linking a word to its’ conceptual representation, and *color-association chunks*, linking a color to its’ conceptual representation. The idea is similar to lemmas from the Altmann model, but in this case both types of dimension-associated chunks need to be retrieved. The Lovett model accounts for individual differences by supporting various strategies to complete the task. In contrast to the Altmann model, this model allows for processing of either stimulus dimension first, but is highly biased towards the word dimension. From either path, chunks associated with the processed dimension are retrieved. Processing can maintain with the retrieval of an answer directly, or the task is checked. Answering directly allows for incorrect answers on incongruent trials and fast responses on congruent trials. When the task is checked, the model compares the dimension of the processed chunk to the goal, which for our purpose is to always respond according to the color of the stimulus. If there is a mismatch, processing continues with the alternative stimulus dimension. Now when retrieving the alternative dimension-associated chunk, the previously retrieved chunk has the same effect as in the Altmann model, facilitating retrieval on congruent trials, having no effect on neutral trials, and inhibiting retrieval on incongruent trials. Notably, this does not necessarily happen on every trial, as there are alternate pathways and strategies, and the model will not retrieve the wrong answer at this point. The base-level activations are set in such a way that incongruent chunks slow retrieval of the correct chunk, and congruent chunks facilitate retrieval of the correct chunk. Once the correct dimension-associated chunk is retrieved, a manual answer is retrieved using the matching chunk, and used to press a key on the keyboard.

The two models offer an ideal comparison for several reasons. First, they deal with an experimental paradigm that is representative of research in cognitive neuroscience. Second, although they embody different and opposing views about the nature of Stroop interference, they are equally successful at predicting the canonical response time effects in the Stroop task (Lovett, 2005; Altmann & Davidson, 2001). Most importantly, these two models exemplify the limits of model identification using behavioral and fMRI data. The two models make use of the same five buffers (visual, motor, goal, imaginal, retrieval). When considering the time needed for perceptual and visual processes (identical in the two models), the difference between the two models is concentrated in a 300 ms window in which different interactions between imaginal, goal, and retrieval buffers are posited. Because the BOLD responses recorded in fMRI are much more sluggish and extend for multiple seconds after a point event, it is reasonable to assume that the two models would make almost identical neuroimaging predictions.

To confirm this suspicion, ACT-R’s canonical BOLD-response prediction tools were used to simulate the neuroimaging responses for the the various experimental conditions in the two models. Fig 2 illustrate the case for incongruent trials. For the sake of illustration, the amplitudes of the BOLD curves were fit so that they would have the same height^1^. It is immediately apparent that the different inter-module dynamics of the two models are lost in the neuroimaging data; all the BOLD curves for all modules are largely overlapping within and between models.

**Figure 2:**
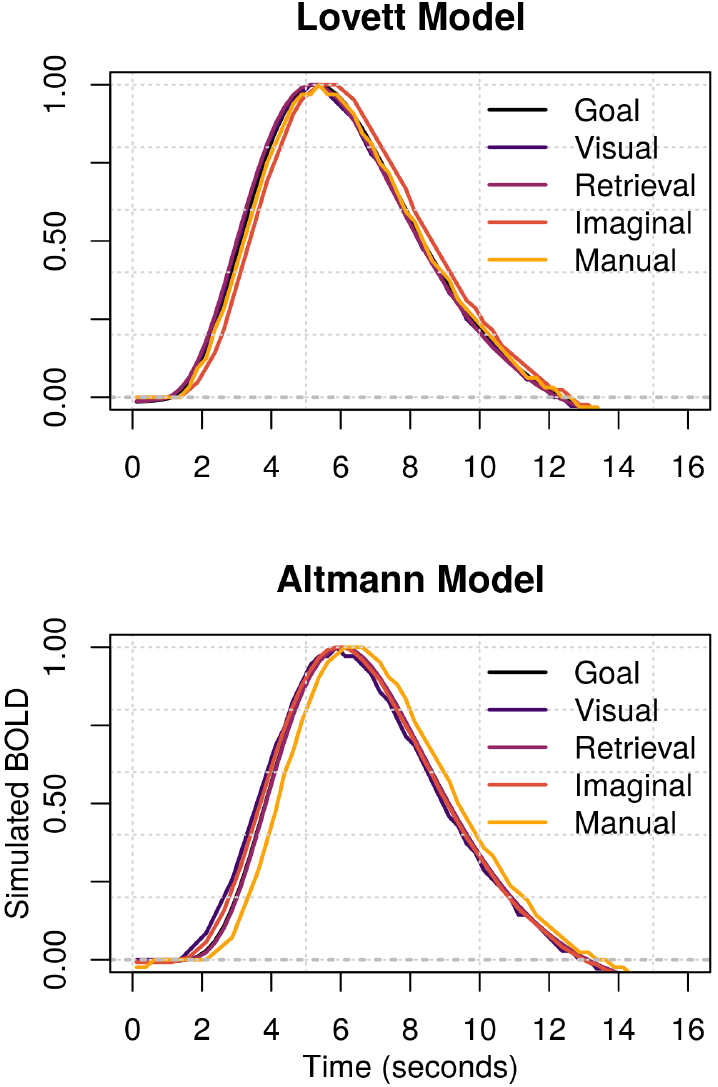
Normalized BOLD-response predictions for incongruent Stroop trials across five different modules in the Lovett (top) and Altmann (bottom) models (See Fig. 1)

Crucially, although these different interactions produce indistinguishable BOLD traces, they *do* produce different *effective connectivity* matrices. And, as the next sections will show, these matrices *do* provide evidence in favor of one model over the other.

## Materials and Methods

### Experimental Dataset

In this analysis, we used fMRI data publicly available from an open repository^2^. The original data was collected at Carnegie Mellon University and published by Verstynen (2014).

### Participants

The dataset contained data from *N* = 30 participants (10 female), aged 21–45 (mean 31). The recruitment procedures can be found in the original publication (Verstynen, 2014).

### Experimental Task

Participants performed a manual-response version of the Stroop task (Stroop, 1935), during which the subjects were asked to indicate the color of a written word presented in the center of the screen. Stimuli could be congruent (“RED” printed in red), incongruent (“RED” printed in green), or neutral (“CHAIR” printed in red). Participants responded by indicating the colors red, green, and blue using the right index, middle, and ring fingers, respectively. Each session consisted of 120 trials (42 congruent, 42 neutral, 36 incongruent) in randomized order.

### Image Acquisition and Preprocessing

As described in Verstynen (2014), the original raw data was acquired using a Siemens Verio 3T system in the Scientific Imaging and Brain Research (SIBR) Center at Carnegie Mellon University with a 32-channel head coil. Functional images were collected using gradient echoplanar pulse sequence with TR = 1,500 ms, TE = 20 ms, and a 90 flip angle. Each volume acquisition consisted of 30 axial slices, each of which was 4 mm thick with 0-mm gap and an in-plane resolution of 3.2 × 3.2 mm. A T1-weighted structural image was also acquired for each participant in the same space as the functional images, but consisting of 176 1-mm slice with with an in-plane resolution of 1 × 1 mm.

For the purpose of our analysis, the original raw data was processed in SPM12 (Wellcome Department of Imaging Neuroscience, www.fil.ion.ucl.ac.uk/spm) following the exactly same preprocessing pipeline as the one indicated in the original publication. Images were corrected for differences in slice acquisition time, spatially realigned to the first image in the series, normalized to the Montreal Neurological Institute (MNI) ICBM 152 template, resampled to 2 × 2 × 2 mm voxels, and finally smoothed with a 8 × 8 × 8-mm full-width-at-half-maximum Gaussian kernel to decrease spatial noise and to accommodate individual differences in anatomy.

### Regions of Interest

DCM analysis is performed on fMRI time-series extracted from specific ROIs. In our case, the ROIs correspond to the specific brain regions that have been previously identified as corresponding to ACT-R buffers. The Talairach coordinates used for each module in the brain followed the convention used by Anderson et al. (2008). The algorithms described in Lacadie, Fulbright, Rajeevan, Constable, and Papademetris (2008) were used to convert Talairach coordinates to Montreal Neurological Imaging Institute (MNI) coordinates. The ROI mask files were created through FSL (Woolrich et al., 2009) of size 16 mm (125 voxels in total) then used to extract fMRI time series from each voxel in each ROI. Principal Component Analysis was then applied on all the extracted time series to identify the time series that best characterized each ROI. The largest principle component was used to project the original data to the new space with more than 75% of the variance explained in each module.

### Dynamic Causal Modeling Analysis

Because DCM is a model-based technique, estimates of connectivity can only derived from parameters corresponding to the specified connectivity between ROIs. To gather complete estimates of connectivity, an unconstrained, fully connected model was generated, in which any ROI was bidirectionally connected to all the others. Furthermore, to identify different patterns of connectivity between conditions, both matrices *B* and *C* were used. Specifically, matrix *C* was used to specify the onset and offset of stimuli, and drive the activity of the “visual” ROI, thus initiating trial-specific activity in the network. In addition, we used the modulatory matrix *B* to specify modulatory effects of condition-specific trials (congruent, neutral, and incongruent) and the ROI connectivity parameters ***A***. Thus, the effective connectivity matrix *E*_*k*_ specific to task condition *k* can be expressed as the element-wise product of ***A*** and the modulatory effects of condition *k* **B**_*k*_, namely:

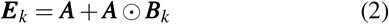

As it is common in DCM, all the parameters were identified using an Expectation-Maximization procedure.

## Results

Figure 3 illustrates the results of the effective connectivity analysis of the fMRI data and the corresponding model predictions. In the figure, columns correspond to the three experimental conditions of the Stroop task (congruent, incongruent, and neutral trials), while the rows correspond to either the predictions of the models (Lovett model, top row; Altmann model, middle row) or the empirical data (bottom row). The reported values of effective connectivity were generated by performing a Bayesian parameter averaging procedure (Kasess et al., 2010) over the individual connectivity matrices generated for each individual participant. Because, in DCM, self-connectivity values need to be set to negative values to ensure the stability of the dynamic state equation (1), the corresponding values were ignored in the analysis and set to zero in Figure 3. Note that the reason we chose Frobenius norm instead of correlation as the metric is that we are interested in the **absolute** measurement of the effective connectivity, not the relative scale between modules. For example, two connectivity vectors of [1, 1, 1, 2, 1] and [−2, −2, −2, −1, −2] would have perfect correlation (*r* = 1), yet they represent opposite connectivity effects (excitatory vs. inhibitory) in all modules. The scale of the values between real fMRI data and ACT-R models may be different, but since all ACT-R models are on the same scale, the differences are still comparable across models.

**Figure 3:**
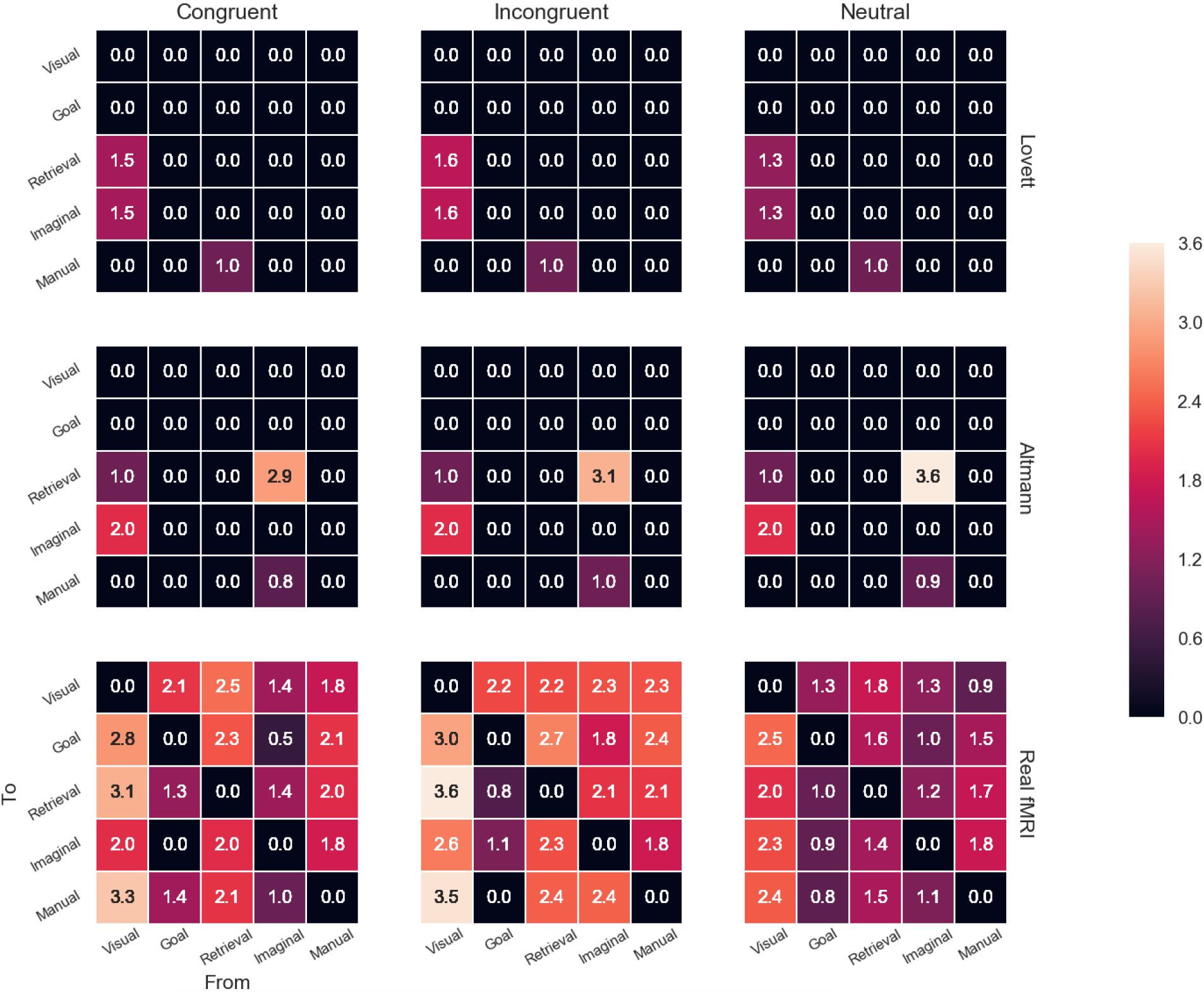
Effective connectivity analysis of the fMRI data and corresponding model predictions (rows), divided by experimental condition (columns).

In general, the connectivity patterns predicted by the two models are much less rich and interconnected than what was measured in the data (Figure 3). This is not unexpected, given the high level of neural abstraction that characterizes ACT-R models (Figure 1). Critically, and as expected, the two models do make different predictions in terms of effective connectivity. To compare the degree of similarity between each model’s predictions and the data, we calculated the Frobenius distance of the difference between the predicted (***P***) and the empirical data matrix (***D***) for each condition *k*, i.e. ||***P***_*k*_ – ***D***_*k*_||_*F*_. This measure can be interpreted as a dissimilarity metric; the smaller the difference between two matrices, the smaller the norm. The results of these comparisons are shown in Figure 4. As shown, the Lovett model yields consistently smaller norm values, and is therefore more similar to the data, across all three conditions.

**Figure 4:**
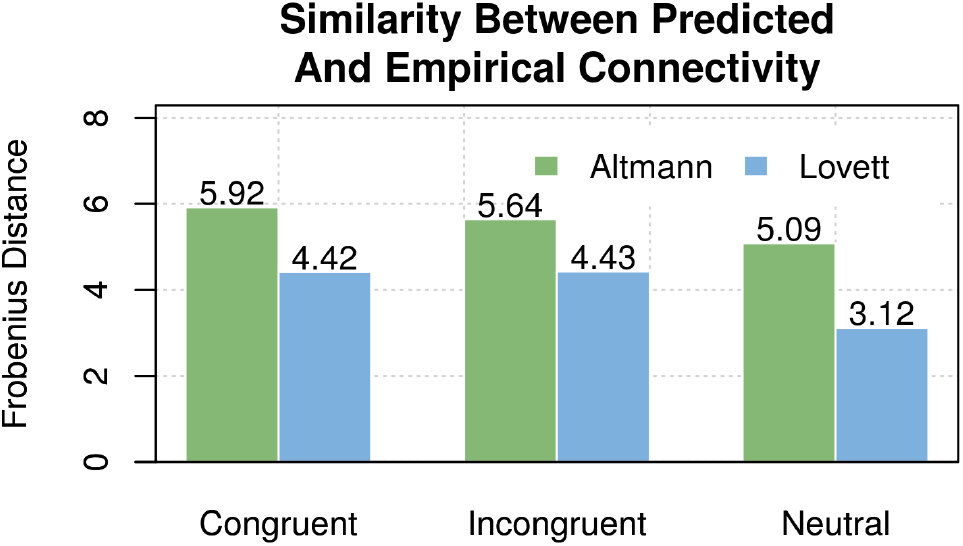
Similarity (Frobenius distance) bewteen predicted and empirical effective connectivity for the Altmann and the Lovett models.

## Discussion

This paper has provided a proof of concept of how analysis of effective connectivity can be used to supplement traditional, GLM-based analysis of neuroimaging data in distinguishing between alternative models. While effective connectivity analysis has been used in cognitive neuroscience for more than a decade, this is the first time, to the best of our knowledge, that this method is used in conjunction with a cognitive modeling approach, and with cognitive architectures in particular. In outlining our method, we choose ACT-R as a modeling paradigm and DCM as a technique to estimate effective connectivity. Neither of these choices, however, are absolute requirements. Connectivity estimates can be gathered from many types of models; the procedure described in this paper certainly applies to other production system-based architecture, like Soar and EPIC, as well. Similarly, although connectivity was estimated with DCM, other methods could be possibly used. For example, Granger Causality. Thus, although we made specific implementation choices, our methods could be instatiated in multiple ways.

Despite encouraging results, a number of limitations need to be acknowledged. First, our method for deriving effective connectivity predictions from ACT-R models is still preliminary. While we believe that it is reasonable, other procedures could be envisioned. For example, operations such as buffer status checks and buffer harvesting could be included in generating our matrices. It is plausible that richer prediction schemes could lead to more realistic connectivity matrices that the ones in Figure 3. It is also plausible that better similarity metrics than Frobenius distance could be used to compare predictions.

These limitations notwithstanding, we see our method as having potential for future modeling research. In particular, we believe that the connectivity matrices obtained from the data can be used to inform model development as well as for model comparison. It is apparent that neither the Lovett nor the Altmann model provide good fits to the data. Because the differences correspond to variables in production rules, the comparison suggests which other production rules or variable bindings could be taking place in the model. In theory, and provided reasonable task constraints, an analysis of the effective connectivity matrices might be used to automatically generate production rules that would match the data. We see this an exciting opportunity for future research.

## Footnotes

1 The amplitude of the BOLD response is a free parameter that can be separately fit for every module; thus, our procedure does not lose generality

2 The data is available on OpenNeuro at the following URL: https://openneuro.org/datasets/ds000164/versions/00001

## References

Altmann, E. M., & Davidson, D. J. (2001). An integrative approach to stroop: Combining a language model and a unified cognitive theory. In Proceedings of the 23rd Annual Meeting of the Cognitive Science Society (pp. 21–26).

Anderson, J. R. (2007). How can the human mind occur in the physical universe? Oxford University Press.

Anderson, J. R., Bothell, D., Byrne, M. D., Douglass, S., Lebiere, C., & Qin, Y. (2004). An integrated theory of the mind. Psychological Review, 111(4), 1036–1060.

Anderson, J. R., Fincham, J. M., Qin, Y., & Stocco, A. (2008). A central circuit of the mind. Trends in Cognitive Sciences, 12(4), 136–143.

Borst, J. P., Nijboer, M., Taatgen, N. A., van Rijn, H., & Anderson, J. R. (2015). Using data-driven model-brain mappings to constrain formal models of cognition. PLoS One, 10(3), e0119673.

Bugg, J. M., McDaniel, M. A., Scullin, M. K., & Braver, T. S. (2011). Revealing list-level control in the stroop task by uncovering its benefits and a cost. Journal of Experimental Psychology: Human Perception and Performance, 37(5), 1595.

Danker, J. F., Gunn, P., & Anderson, J. R. (2008). A rational account of memory predicts left prefrontal activation during controlled retrieval. Cerebral Cortex, 18(11), 2674–2685.

Fincham, J. M., & Anderson, J. R. (2006). Distinct roles of the anterior cingulate and prefrontal cortex in the acquisition and performance of a cognitive skill. Proceedings of the National Academy of Sciences, 103(34), 12941–12946.

Friston, K., Harrison, L., & Penny, W. (2003). Dynamic Causal Modelling. NeuroImage, 19(4), 1273–1302.

Kasess, C. H., Stephan, K. E., Weissenbacher, A., Pezawas, L., Moser, E., & Windischberger, C. (2010, feb). Multi-subject analyses with dynamic causal modeling. NeuroImage, 49(4), 3065–3074.

Kotseruba, I., & Tsotsos, J. K. (2018). 40 years of cognitive architectures: core cognitive abilities and practical applications. Artificial Intelligence Review, 1–78.

Lacadie, C. M., Fulbright, R. K., Rajeevan, N., Constable, R. T., & Papademetris, X. (2008, aug). More accurate talairach coordinates for neuroimaging using non-linear registration. NeuroImage, 42(2), 717–725.

Lovett, M. C. (2005). A strategy-based interpretation of stroop. Cognitive Science, 29(3), 493–524.

Pitt, M. A., Kim, W., Navarro, D. J., & Myung, J. I. (2006). Global model analysis by parameter space partitioning. Psychological Review, 113(1), 57.

Rice, P., & Stocco, A. (2019). The role of dorsal premotor cortex in resolving abstract motor rules: Converging evidence from Transcranial Magnetic Stimulation and cognitive modeling. Topics in cognitive science, 11(1), 240–260.

Sohn, M.-H., Albert, M. V., Jung, K., Carter, C. S., & Anderson, J. R. (2007). Anticipation of conflict monitoring in the anterior cingulate cortex and the prefrontal cortex. Proceedings of the National Academy of Sciences, 104(25), 10330–10334.

Sohn, M.-H., Goode, A., Koedinger, K. R., Stenger, V. A., Fissell, K., Carter, C. S., & Anderson, J. R. (2004). Behavioral equivalence, but not neural equivalenceneural evidence of alternative strategies in mathematical thinking. Nature Neuroscience, 7(11), 1193.

Stocco, A., Lebiere, C., & Anderson, J. R. (2010). Conditional routing of information to the cortex: A model of the basal ganglias role in cognitive coordination. Psychological Review, 117(2), 541.

Stroop, J. R. (1935). Studies of interference in serial verbal reactions. Journal of Experimental Psychology, 18(6), 643–662.

van Vugt, M. K. (2014). Cognitive architectures as a tool for investigating the role of oscillatory power and coherence in cognition. NeuroImage, 85, 685–693.

Verstynen, T. D. (2014, nov). The organization and dynamics of corticostriatal pathways link the medial orbitofrontal cortex to future behavioral responses. Journal of Neurophysiology, 112(10), 2457–2469.

Woolrich, M. W., Jbabdi, S., Patenaude, B., Chappell, M., Makni, S., Behrens, T., … Smith, S. M. (2009, mar). Bayesian analysis of neuroimaging data in FSL. NeuroImage, 45(1), S173–S186.

